# Increased listening effort and decreased speech discrimination at high presentation sound levels in acoustic hearing listeners and cochlear implant users

**DOI:** 10.1101/2024.09.20.614145

**Authors:** Chengjie G. Huang, Natalie A. Field, Marie-Elise Latorre, Samira Anderson, Matthew J. Goupell

## Abstract

The sounds we experience in our everyday communication can vary greatly in terms of level and background noise depending on the environment. Paradoxically, increasing the sound intensity may lead to worsened speech understanding, especially in noise. This is known as the “Rollover” phenomenon. There have been limited studies on rollover and how it is experienced differentially across aging groups, for those with and without hearing loss, as well as cochlear implant (CI) users. There is also mounting evidence that listening effort plays an important role in challenging listening conditions and can be directly quantified with objective measures such as pupil dilation. We found that listening effort was modulated by sound level and that rollover occurred primarily in the presence of background noise. The effect on listening effort was exacerbated by age and hearing loss in acoustic listeners, with greatest effect in older listeners with hearing loss, while there was no effect in CI users. The age- and hearing-dependent effects of rollover highlight the potential negative impact of amplification to high sound levels and therefore has implications for effective treatment of age-related hearing loss.

## Introduction

Auditory processing of speech is critical to social communication and interaction in everyday life and is complicated by dynamic factors, influencing a person’ ability to perceive speech in any given environment. For example, speech perception difficulties occur more frequently in noisy environments with background noise or multiple talkers than in a quiet environment without any noise (Dubno et al., 1984; Freyman et al., 2004; Banh et al., 2012). Speech-perception difficulties are exacerbated by hearing loss, and hearing aids or cochlear implants (CIs) often fail to improve speech understanding in noise for hearing-impaired (HI) and CI users. Paradoxically, increasing the sound intensity of speech and noise to increase audibility does not necessarily yield better speech understanding, and instead higher sound intensities actually decrease performance, especially in noise (Molis and Summers, 2003; Summers and Cord, 2007). Therefore, such amplification provided for HI listeners does not necessarily remedy their poor speech-in-noise understanding. Interestingly, this effect has been found in both normal-hearing (NH) and HI individuals, as well as CI users. This phenomenon is known as “rollover”, and it is poorly understood in general.

Clinically, rollover is defined as a reduction in speech recognition scores that occurs at intensities above the level where performance is at a maximum. Clinical rollover is traditionally used as a diagnostic tool for the indication of retrocochlear pathologies, such as a vestibular schwannoma (Jerger and Jerger, 1971; Dirks et al., 1977). Note that these studies are performed in the absence of background noise. In addition, this reduction must surpass a criterion to be considered clinically relevant and may not capture the full extent of speech perception difficulties in individuals who show no evidence of clinical pathology. In contrast to the clinical diagnosis of rollover, more recent studies of rollover demonstrate that it is not limited to cases of retrocochlear pathologies (Studebaker et al., 1999; Molis and Summers, 2003; Hornsby et al., 2005). These studies were performed on adults with audiometrically normal hearing thresholds, suggesting that rollover is also present in the absence of retrocochlear pathologies, and exacerbated by the presence of noise or degraded speech.

Studies concerning the rollover phenomenon have yielded limited success in generating hypotheses for the underlying mechanisms. Aging affects speech perception, and many older individuals complain of difficulties in speech understanding in noise (Middelweerd et al., 1990; Banh et al., 2012; Tremblay et al., 2015; Pang et al., 2019), despite having similar clinical audiograms as younger individuals, and thus are classified as “normal hearing.” This problem is compounded by issues related to age-related hearing loss, where hearing status interacting with aging may contribute to rollover. The intersection between the effects of aging and poor speech understanding in noise, and how this relates to listening at high sound intensities is also unclear and remains a central problem in auditory neuroscience. Furthermore, it is unclear whether rollover is caused by a spectral or temporal deficit, or a combination of both as there have been limited and conflicting results in the literature (Molis and Summers, 2003; Hornsby et al., 2005; Hornsby and Ricketts, 2006; Summers and Cord, 2007). These hypotheses must therefore be systematically tested to elucidate the interaction between aging, hearing loss and the resulting spectrotemporal deficits.

Another dimension of speech perception besides performance is the effort required to achieve that performance. Listening in noisy environments requires “deliberate allocation of mental resources”, which may include several facets of effortful listening including increased arousal, attention, working memory, and auditory processing (Pichora-Fuller et al., 2016). Objective measurement of listening is commonly done with pupillometry (Winn et al., 2018). Because pupil dilation is an unconscious biophysical change reflecting cognitive effort, difficult listening conditions associated with greater listening effort led to an increase in pupil dilation (Zekveld et al., 2010; Winn et al., 2015). Increased listening effort can compensate for difficult listening environments with advancing age as well as hearing status (Ayasse et al., 2016; Ayasse and Wingfield, 2018). However, there is an upper limit to the extent to which increased effort can improve speech understanding in HI or CI listeners (Hornsby, 2013; Bess and Hornsby, 2014). Assessment of speech perception alone might not fully reveal the mechanisms underlying rollover, since effort can be modulated to achieve similar levels of behavioral performance. Therefore, using pupil dilation to assess how aging and hearing status interact with rollover of speech is a novel approach that allows us to decouple effort contributions for the resulting behavioral perception at high sound intensities. The insights gained from this study will reveal how top-down modulatory processes such as listening effort affect performance in difficult listening situations that are worsened by rollover.

Here we used a speech discrimination task and pupillometry to assess the extent to which younger and older NH (YNH and ONH respectively), older HI (OHI), and older CI (OCI) experience rollover as well as the deployment of listening effort at high presentation levels. Minimal word pairs were presented in both quiet and with background 6-talker babble noise across multiple sound intensities to obtain a comprehensive understanding of rollover. CI users often experience some of the same problems as HI listeners, with degraded speech perception due to poor spectral resolution, as well as other highly varied problems including electrode-to-neural interface, which may directly affect auditory information processing (Zhou et al., 2019; Shader et al., 2020; Johnson et al., 2021). Therefore, CI users may help us untangle the mechanisms which underlie rollover. We hypothesized that rollover is experienced differentially between groups and that aging and hearing loss exacerbates the effects of rollover. In addition, we hypothesized that listening effort is similarly modulated by high sound intensities, where different groups will expend more effort when sound level exceeds past a comfortable hearing level (≥65 dB SPL). Specifically, we hypothesized that ONH listeners would expend increased effort compared to YNH listeners to offset rollover, in that more effort is needed to compensate as individuals age. These compensatory mechanisms are limited by hearing impairment in HI and CI listeners, and thus we hypothesized that despite increased listening effort, they cannot fully compensate for speech perception deficits at higher levels compared with NH listeners.

## Materials and Methods

### Listeners

Native-English speakers were recruited for this study for each of the following groups: young NH (YNH, 21-25 years, N = 14), middle-aged to older NH (ONH, 52-76 years, N = 17), middle-aged to older HI (OHI, 53-81 years, N = 16), and middle-aged to older CI users (OCI, 55-84 years, N = 20). Normal hearing was defined as pure-tone thresholds ≤ 25 dB HL (re: ANSI 2018) at each frequency tested from 250 to 4000 Hz in both ears. Hearing impairment was defined as pure-tone thresholds > 25 dB HL and < 65 dB HL at each frequency tested from 250 to 4000 Hz in both ears, as our criteria only included listeners with “mild-to-moderate” hearing loss. Additional criteria for all listeners included the following: A passing score of ≥ 26 on the Montreal Cognitive Assessment (Nasreddine et al., 2005) or HI-MoCA (Dawes et al., 2019), normal or corrected-to-normal vision, and a negative history of neurological disease, middle ear surgery, or untreated vision issues. All procedures were reviewed and approved by the Institutional Review Board at the University of Maryland, College Park. Listeners provided informed consent and were compensated for their time.

### Equipment

Listeners were seated in a double-walled sound-attenuating booth (IAC Acoustics, North Aurora, IL) in front of a desktop computer where they performed the task. The auditory stimuli were presented to both ears of listeners through circumaural headphones (Sennheiser HD650, Hanover, Germany) while they viewed a computer monitor. For CI listeners, headphones were placed over the behind-the-ear processors (Ricketts et al., 2006; Goupell et al., 2018). Visual and auditory stimulus presentations were controlled using custom E-Prime scripts (Psychology Software Tools, Pittsburgh, PA) and amplified using a custom sound card routed through a Chronos box (Psychology Software Tools, Pittsburgh, PA). Behavioral responses were recorded online to a network drive connected to the desktop computer that was also used by the listener to enter responses via a keyboard. Pupil data were collected using a desktop-mounted Eyelink 1000 Plus Monocular system (SR Research, Ottawa, Canada) at a sampling rate of 1000 Hz. Pupil tracking was calibrated and validated with a nine-point grid at the start of each run, and monocular tracking was used to monitor either left or right eye gaze and pupil size. Listeners were seated approximately 26 in. away from a 24 in. monitor with their chin placed on a chin rest. Testing was completed in one 3-h session, with breaks given as needed.

### Stimuli

The stimuli consisted of eight possible minimal word pairs including four temporal contrast (Gordon-Salant et al., 2006) word pairs (beak-peak, beat-wheat, wheat-weed, dish-ditch) and four non-temporal contrast word pairs (hall-wall, blood-blush, chin-shin, lamb-ram). The word pairs were presented in randomized order across 32 trials per block in either a quiet condition or a noise condition containing 6-talker babble presented at 0-dB SNR at a specific sound intensity (8 word-pairs × 2 orders × 2 listening conditions) in each block. These 32-trial blocks were presented at 35, 55, 65, 75, and 85 dB SPL. Each participant listened to at least two runs of the experiment (i.e., at least 10 randomized blocks).

#### Procedure

Each session began with calibration and validation of the pupil position to obtain baseline coordinates for either the left or right eye. This was followed by a blank screen on the monitor which changed from black to gray to white across 135 s, with 45 s on each screen. This procedure was used to obtain the dynamic range of the pupil to control for individual differences in pupil diameter (Piquado et al., 2010). Participants were instructed to look at the center of the screen and fixate on a red cross to prevent eye drift or errors in tracking pupil dilation across the dynamic range measurement. The average pupil size during the black screen gave an estimate of maximum pupil dilation, and the average pupil size during the white screen gave an estimate of minimum pupil dilation. The estimates from each test session were used as the dynamic range for the data from that session.

The experiment then initiated with the auditory presentation (at t = 500 ms) of a minimal-word pair presented at the chosen stimulus sound intensity while the participant fixated on the same red cross described above on a gray background. After the auditory presentation of the stimulus, the red cross would change to green and both words in the minimal word-pair would appear on the screen (at t = 4500 ms), with each word appearing on either the left side or the right side of the screen. Participants were instructed to indicate on which side of the screen the second word they heard was located. Words presented on the screen were displayed in Courier New monospaced size 72 pt font. For example, if the word-pair presented through the headphones was “beak-peak”, the second word would be “peak” and if “peak” appeared on the left side of the screen, the participant was instructed to press “1” on the keyboard, and the converse would be true if “peak” appeared on the right side of the screen in which case the participant was instructed to hit “2” on the keyboard. The next trial would then begin 6000 ms after their keyboard press to allow pupil area to return to baseline. The trials repeated until all 32 possible pairs were presented across conditions for a specific sound intensity, and the whole block was repeated across the five sound intensities across at least two runs as described above. Figure 1 shows a summary of the experimental procedure.

**Figure 1.**
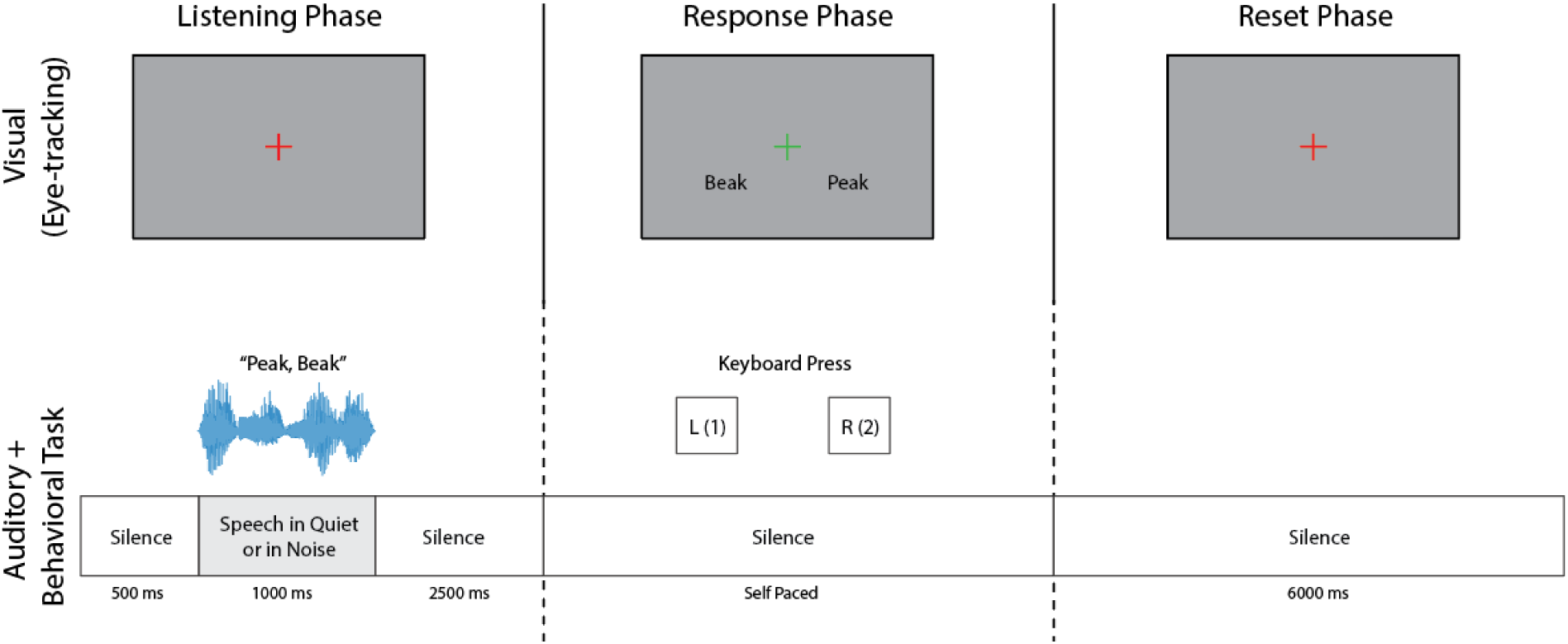
Schematic of experimental paradigm. Speech perception task initializes with a listening phase (left) of 4000 ms where minimal word pairs are presented via headphones after an initial 500 ms of silence. Listeners fixate on a grey screen with a red cross during this phase. The response phase (middle) begins when the red cross turns green and the two words presented are shown on the screen. Participants either press 1 or 2 to indicate which word was heard second in the listening phase. The reset phase (right) occurs after the participant provides a keyboard press lasting 6000 ms to allow pupils to return to baseline.

### Behavioral analysis

Each trial of the experiment was marked as “correct” or “incorrect” depending on if the participant’s keyboard press matched the position of the second word they had heard in the word-pair presented. Behavioral performance was assessed as a score out of 16 for each of the quiet and noise conditions and transformed into a value of percentage correct. The percentage correct was then plotted across sound intensities from 35-85 dB SPL and the amount of rollover was assessed by comparing the behavioral performance at the peak, which usually occurred around a comfortable hearing level and the behavioral performance at the lowest level, which usually occurred at a sound intensity higher than a comfortable listening level. Additional statistical analysis was performed using these data, which is described in further detail below.

### Pupillometry analysis

Preprocessing of eye tracking data was performed similarly to previous pupillometry experiments (Piquado et al., 2010; Zekveld et al., 2010, 2011; Kuchinsky et al., 2013; Kuchinsky et al., 2014) and as recommended for measuring listening effort (Winn et al., 2018). An 11-s window of pupil size over time was preprocessed. Blinks and saccades were removed and replaced with linear interpolations at a threshold of 50%. This means that if more than half of the baseline was interpolated, the trial was removed from analysis. In addition, if more than half of the entire trial was interpolated, the trial was removed from and not considered for data analysis. For the included trials, a five-point moving window was used to smooth each pupil recording, and the data were downsampled to 100 Hz. The pupil area was then transformed into a percentage of the total dynamic range, which was obtained using the average values collected for maximum and minimum pupil dilation on the initial black and white screens described above. The sound onset begins at t = 500 ms after the initial baseline pupil area has been collected. After sound offset, an additional 2500 ms of silence occurs before the response phase begins. Therefore, in pupillometry plots, sound onset is defined as t = 500 ms and response onset is defined as t = 4500 ms.

## Statistical analysis

### Behavior

For the behavior data, we first performed a 3-way omnibus mixed analysis of variance (ANOVA) with factors of group (YNH, ONH, OHI, OCI), SNR (quiet vs. noise), and level (35, 55, 65, 75, 85 dB SPL). Data in percentage of correct words was transformed to rationalized arcsine units (RAUs) in order to account for violations of homogeneity (Studebaker, 1985). For this and subsequent analyses, when there was a violation of the assumption of sphericity, a Greenhouse-Geisser correction was applied.

Due to the significant interaction between the three variables, we separated the data into the quiet condition and the noise condition for further analyses. Within each SNR, we performed mixed ANOVAs with factors of level and group and level x group interaction. Once each of those effects were established, we then performed planned one-way repeated-measures ANOVAs and corrected and adjusted the p-values with Bonferroni method for multiple comparisons. This allowed us to test our hypothesis concerning rollover, which suggests non-monotonic performance as a function of level and a peak in performance. In other words, the key comparisons were between the different levels to find the peak performance.

### Pupillometry

For the pupillometry data, our primary goal was to identify time intervals where the pupil area differed between the quiet and noise conditions. To identify these time intervals, we utilized a nonparametric bootstrap-based statistical analysis (Efron and Tibshirani, 1993; Contadini-Wright et al., 2023). We treated each pupil area trace as a time series for each participant and we bootstrapped trials of each block at each sound level separately. The bootstrapped trials were iterated for 1000 repetitions with replacement and each time point in the traces were compared in an A-B manner. If the proportion of bootstrapped iterations fell above or below zero was 95% (i.e., p = 0.05) of the total iterations, then that given time point would be deemed significant. We performed this analysis for the first 10s from the beginning of the listening phase.

## Results

### Acoustic listeners and CI users experience rollover to different extents in noise

We presented both temporal contrast (Fig. 2A) and non-temporal contrast (Fig. 2B) word pairs both in quiet and in the presence of background babble noise to participants to test whether rollover was present across our groups. We first performed a 4-way mixed ANOVA omnibus test and initially found main effects of SNR [F(1,63) = 374.281, p < 0.001, η_p_^2^ = 0.856], Level [F(4,252) = 26.168, p < 0.001, η_p_^2^ = 0.293], Temporality [F(1,63) = 26.897, p < 0.001, η_p_^2^ = 0.299], and Group [F(3,63) = 63.471, p < 0.001, η_p_2 = 0.751]. Significant interactions of note include SNR x Group [F(3,63) = 28.776, p < 0.001, η_p_^2^ = 0.578], Level x Group [F(12,252) = 4.994, p < 0.001, η_p_^2^ = 0.192], SNR x Level [F(4,252) = 10.108, p < 0.001, η_p_^2^ = 0.138] and a 3-way interaction of SNR x Group x Level [F(12,252) = 4.633, p < 0.001, η_p_^2^ = 0.181]. Temporality had interaction effects with Group [F(3,63) = 6.36, p < 0.001, η_p_^2^ = 0.232], and SNR [F(1,63) = 21.947, p < 0.001, η_p_^2^ = 0.258], but not Level [F(4,252) = 2.257, p = 0.064, η_p_^2^ = 0.035]. Additionally, there were no 3-way or 4-way interactions with Temporality. Furthermore, because there was not a significant interaction of Temporality x Level, we continued the remainder of the analysis without segregating the type of word contrast.

**Figure 2.**
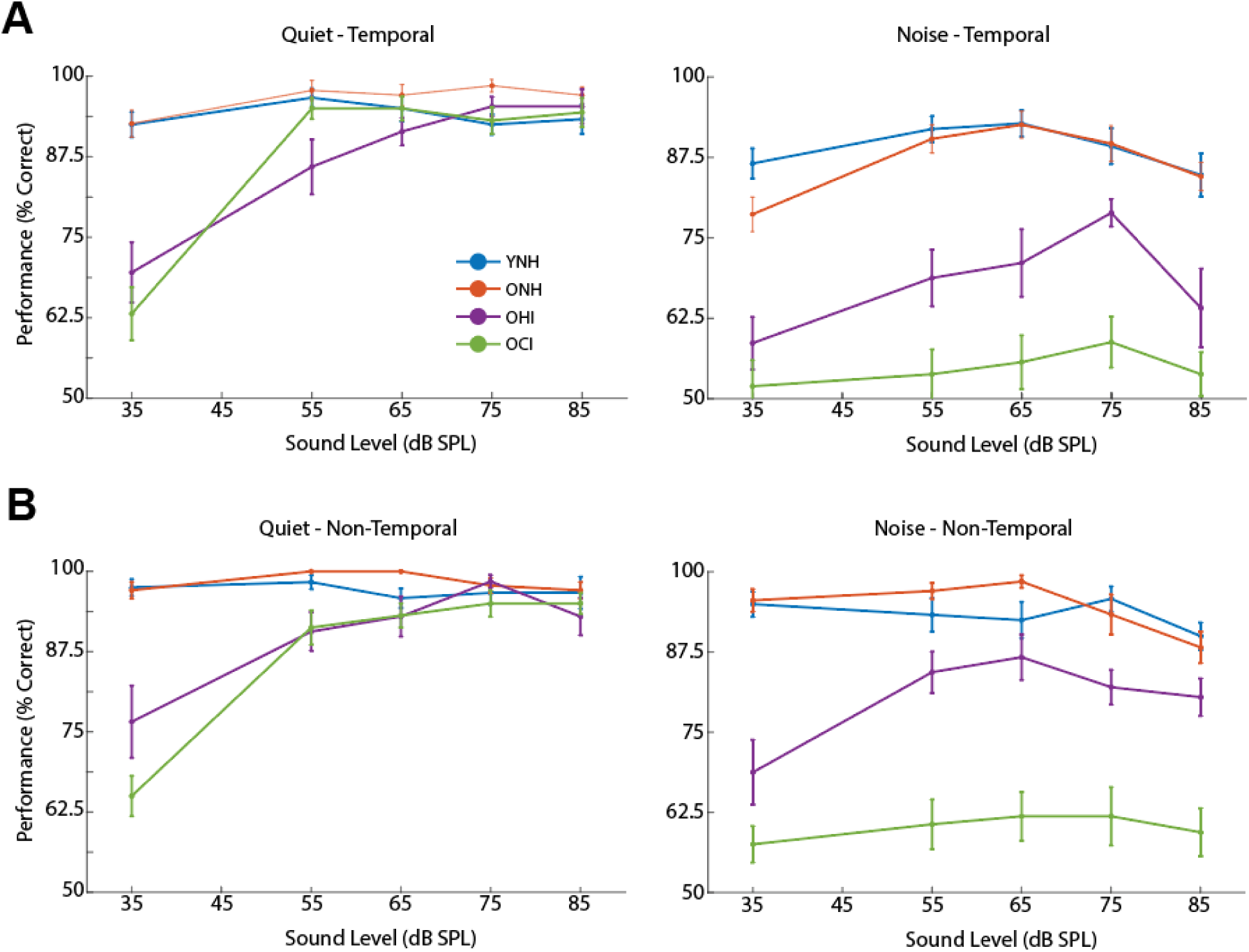
Behavioral performance on experimental task segregated by type of contrast. A) Mean performance of participants for YNH (blue), ONH (orange), OHI (purple) listeners, and OCI (green) users as a function of sound level when speech perception task with temporal contrast word pairs presented in quiet (left) and in noise (right). B) Same as A but for task performed with non-temporal contrast word pairs. Error bars indicate ± 1 standard error.

Combining all the word pairs into our statistical analysis and using a three-way mixed ANOVA, we found main effects of SNR [F(1,60) = 260.845, p < 0.001, η_p_^2^ = 0.813], Group [F(3,60) = 58.779, p < 0.001, η_p_^2^ = 0.746], Level [F(4,240) = 35.152, p < 0.001, η_p_^2^ = 0.369]. The significant effect of SNR is primarily due to worse performance in the noise condition compared to in the quiet condition. In addition, we found significant interactions of SNR x Group [F(3,60) = 23.966, p < 0.001, η_p_^2^ = 0.545], SNR x Level [F(3.3,196.3) = 5.511, p < 0.001, η_p_^2^ = 0.084], and Level x Group [F(12,240) = 3.994, p < 0.001, η_p_^2^ = 0.166]. Finally, we found a significant 3-way interaction of SNR x Group x Level [F(9.8,196.3) = 4.04, p < 0.001, η_p_^2^ = 0.168].

To understand the three-way significant interaction, we decided to separately investigate the quiet and noise conditions. Using a two-way mixed ANOVA, in the quiet condition, our statistics revealed significant main effects of Group [F(3,61) = 18.137, p < 0.001, η_p_^2^ = 0.471] and Level [F(3.5,212.7) = 19.923, p < 0.001, η_p_^2^ = 0.246]. We also found a significant interaction of Level x Group (Two-Way Mixed ANOVA, F[10.5,212.7) = 4.491, p < 0.001, η_p_^2^ = 0.181]. Qualitatively, the YNH and ONH groups performed comparatively near maximum (>90% correct) at all sound intensities. The OHI and CI groups performed near maximum and similarly to the YNH and ONH listeners at 55-85 dB SPL; however, they performed worse at 35- and 55-dB SPL. This was demonstrated quantitatively with our detailed post-hoc statistical analyses (Supplementary Table 1, Fig. 3A).

**Figure 3.**
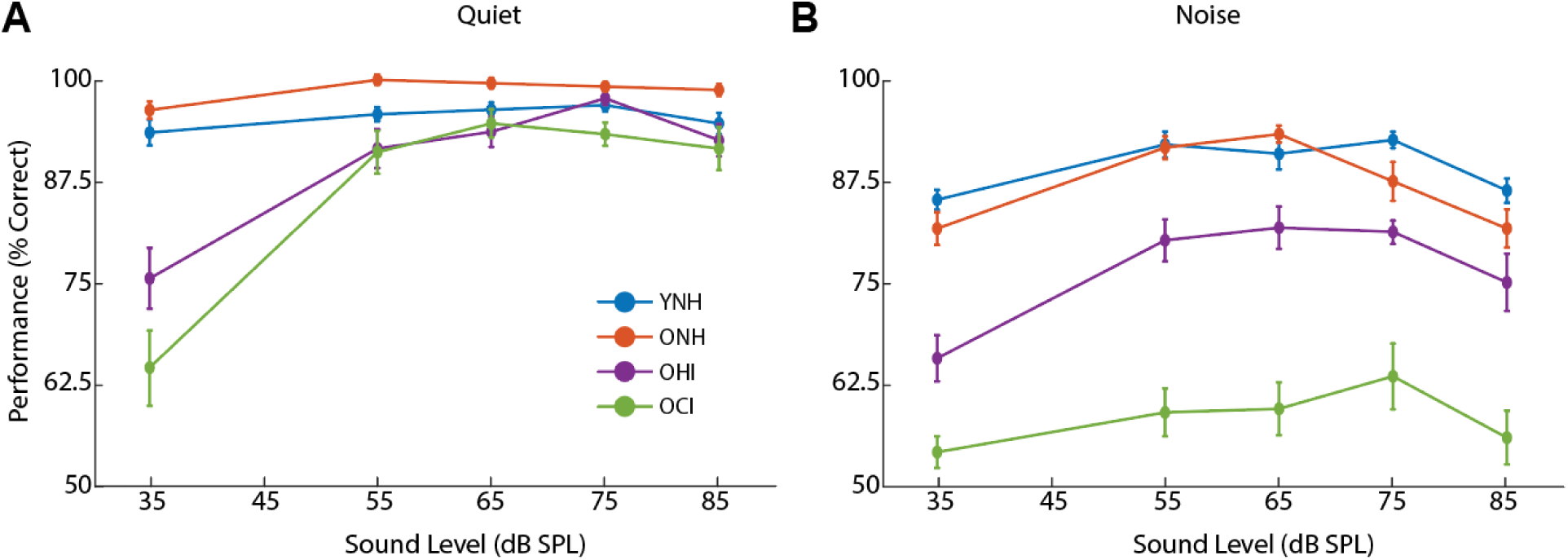
Behavioral performance on experimental task. A) Mean performance of participants for YNH (blue), ONH (orange), OHI (purple) listeners, and OCI (green) users as a function of sound level when speech perception task was performed in quiet. B) Same as A but for task performed in babble noise. Error bars indicate ± 1 standard error. For detailed statistics, refer to Supplementary Table 1 and 2.

To determine if there was significant rollover in the quiet conditions, one-way repeated-measures ANOVAs revealed that there were no main effects of level for the YNH [F(4,44) = 0.042, p = 0.997, η_p_^2^ = 0.004] and ONH [F(4,60) = 2.560, η_p_^2^ = 0.146, p = 0.05, η_p_^2^ = 0.146] groups, but there were for OHI [F(4,60) = 10.400, p < 0.001, η_p_^2^ = 0.409] and CI [F(4,80) = 19.284, p < 0.001, η_p_^2^ = 0.491]. The significance from the OHI and CI groups were largely driven by the mean performances at 35 dB SPL, as there was no significant difference in any of our groups when comparing behavioral performance at a comfortable hearing level at 65 dB SPL and the highest sound level at 85 dB SPL (Supplementary Table 2). This demonstrates that rollover is in fact not present when our stimuli were presented in quiet across all of our testing groups.

In contrast to our results in quiet, when the same speech tokens were presented in 6-talker babble noise, rollover was experienced by all testing groups, and to different extents. Overall, all groups significantly dropped in behavioral performance from the quiet condition across all sound levels. Using a two-way mixed ANOVA, in the noise condition, we found significant main effects of Group [F(3,60) = 74.517, p < 0.001, η_p_^2^ = 0.788] and Level [F(4,240) = 21.313, p < 0.001, η_p_^2^ = 0.262]. We also found significant interactions for Level x Group as we did for our data in the quiet condition [F(12,240) = 3.561, p < 0.001, η_p_^2^ = 0.151].

To determine if there was significant rollover in the noise conditions, one-way repeated-measures ANOVAs revealed that there was no significant difference in performance [F(4,44) = 2.451, p = 0.06, η_p_^2^ = 0.182] for YNHs when comparing across levels (Fig. 3B, blue curve). For the ONH listeners, there was a significant effect of level [F(4,56) = 7.561, p < 0.001, η_p_^2^ = 0.353] and this was driven by a decrease when comparing their performance at 65 with 85 dB SPL (p < 0.001, Fig. 3B, orange curve). For the OHI listeners, there was also a significant effect of level [F(4,60) = 20.327, p < 0.001, η_p_^2^ = 0.575] and this was driven by decrease in performance between 65 and 85 dB SPL (p = 0.023, Fig. 3B, purple curve). For the CI users, overall behavioral performance was lower than all other testing groups at all sound levels, and peaked at 75 dB SPL instead, but was not significantly different across levels (Fig. 3B, green curve). This is important to note since Fig. 3A demonstrates that CI users can perform the task and the drop in performance in Fig. 3B demonstrates that they were not at the noise floor or performing at chance level (50%). All statistical comparisons are summarized in Supplementary Table 2. Combined, these results show that rollover is a perceptual phenomenon that is experienced when stimuli are presented in noise and is absent in quiet. Furthermore, rollover is experienced by acoustic listeners to different extents depending on age and hearing status, independent of audibility, as rollover occurs once audibility is achieved.

### ONH listeners expend more effort than YNH listeners with increased sound levels

One particularly interesting finding from our behavioral results was that YNH and ONH listeners performed similarly to each other in quiet and in noise. In fact, as noted above, YNH, ONH, and OHI listeners experienced significant rollover when comparing their performance at 65 and 85 dB SPL, with a slightly larger decrease in ONH listeners compared to YNH listeners (Fig. 3B, blue and orange curves). To explore this idea further, we wanted to understand whether listening effort played a role in similar behavioral performance between YNH and ONH listeners. In order to assess listening effort, we measured pupil area for each participant while they were performing the speech perception task. We measured pupil area across the entire duration of the trial and focused in particular on the pupil area response after the stimulus onset and the response onset when the red cross on the screen turns green. We calculated the pupil response as a percentage of each participant’s dynamic range to control for variability between testing groups similar to Milvae et al. (2021). At 65 dB SPL, pupil area significantly increased after sound onset when speech stimuli were presented in noise for ONH listeners (1900-3990 ms, p < 0.05, Fig. 4B), but not for YNH listeners (Fig. 4A). At 85 dB SPL, pupil area instead significantly increased after sound onset for both YNH (740-2350 ms) and ONH (2120-3730 ms) listeners in the noise condition (p < 0.05, Fig. 4C, 4D). In particular, the ONH group experienced a significantly larger increase in pupil area compared to the YNH group in the noise condition in addition to a longer duration of significantly increased pupil size compared to the YNH group (compare Fig. 4C and Fig. 4D). Taken together with the behavioral results, this demonstrates that while YNH and ONH listeners can perform the task similarly well while experiencing a similar degree of rollover, the ONH listeners appear to expend more listening effort than YNH listeners at both 65 and 85 dB SPL.

**Figure 4.**
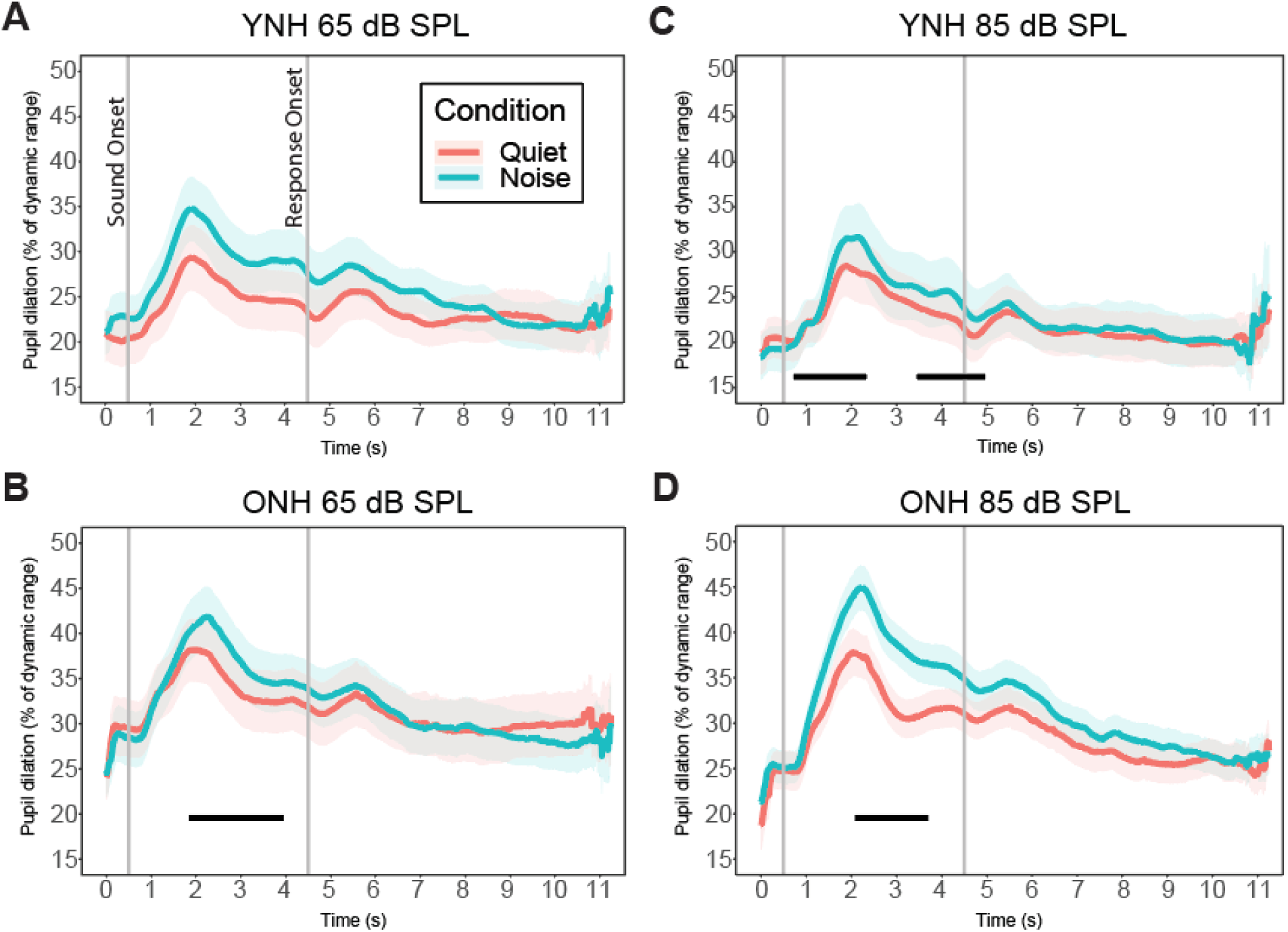
Pupillometry data for normal-hearing listeners. A) Mean pupil dilation as a percentage of dynamic range across the timescale of each trial of experimental task performed in quiet (orange) and in the presence of noise (cyan) at 65 dB SPL for YNH listeners. B) Same as A but for ONH listeners. C and D) Same as A and B respectively but at 85 dB SPL. Shaded error bars indicate ± 1 standard error. Black bars indicate significantly different pupil size between quiet and noise conditions at the p < 0.05 level across the timescale shown.

### OHI listeners demonstrate extended listening effort with increased sound level

For OHI listeners, pupillometry data revealed that there was no significant difference in pupil area between the quiet and noise condition at 65 dB SPL. However, it is important to note that the pupil area baseline occurred at a significantly higher level than the NH listeners at ∼30% of their pupil dynamic range (p < 0.05, Fig. 5A). In contrast, when stimuli were presented at 85 dB SPL, there was a significant increase in pupil area in the noise condition compared to the quiet condition (2880-4710 ms, p < 0.05, Fig. 5B), demonstrating some degree of compensation by listening effort. In addition, the elevated increase in pupil size extended beyond into the response phase between 6020-10000 ms, indicating sustained increased effort even while responding to the task. Despite this increase in listening effort, OHI listeners could not perform behaviorally at a similar level as the NH listeners (Fig. 3B) and thus were unable to compensate insufficiently.

**Figure 5.**
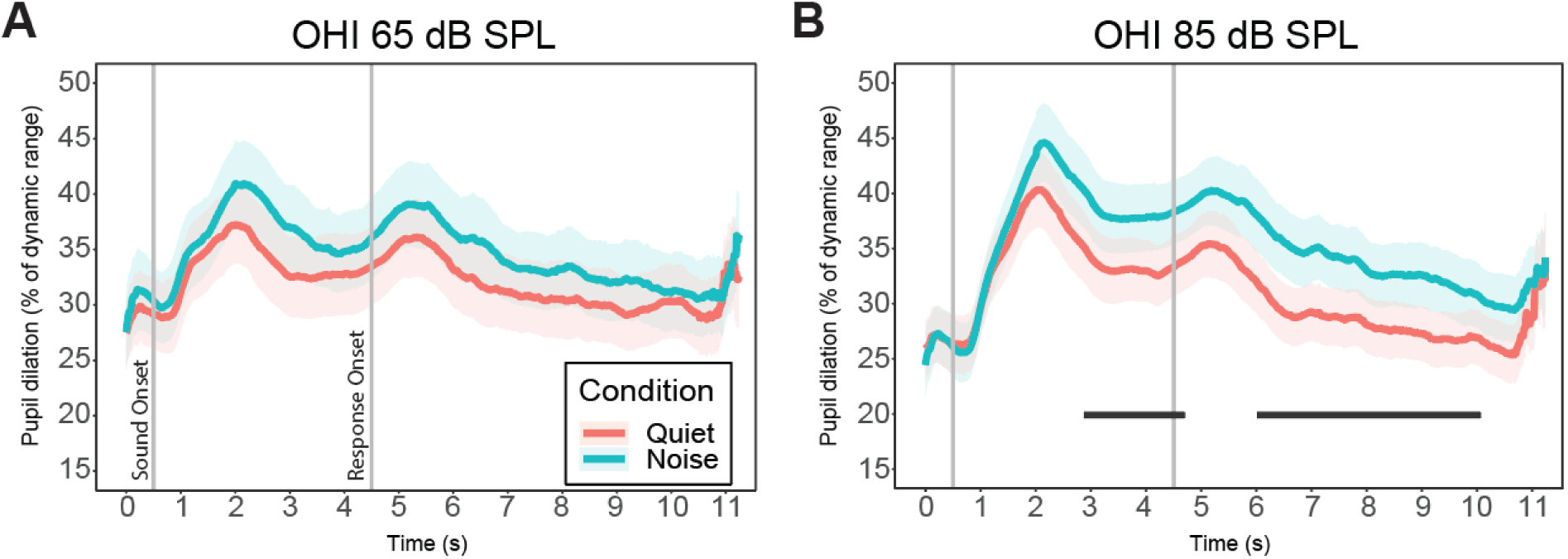
Pupillometry data for OHI listeners. A) Mean pupil dilation as a percentage of dynamic range across the timescale of each trial of experimental task performed in quiet (orange) and in the presence of noise (cyan) at 65 dB SPL for OHI listeners. B) Same as A but at 85 dB SPL. Shaded error bars indicate ± 1 standard error. Black bars indicate significantly different pupil size between quiet and noise conditions at the p < 0.05 level across the timescale shown.

### OCI users display no increased listening effort with increased sound levels

For older CI users, there was a brief significant period of increased pupil size at 6810-7030 ms present in the response phase at 65 dB SPL; however, there was no increased pupil size during the listening phase for either 65 dB or 85 dB SPL (Fig. 6A and 6B). These results demonstrated that there is not increased listening effort when comparing the quiet and noise conditions with increasing sound level. Supplemental figure 1 shows a summary of all of our pupillometry data across listening groups.

**Figure 6.**
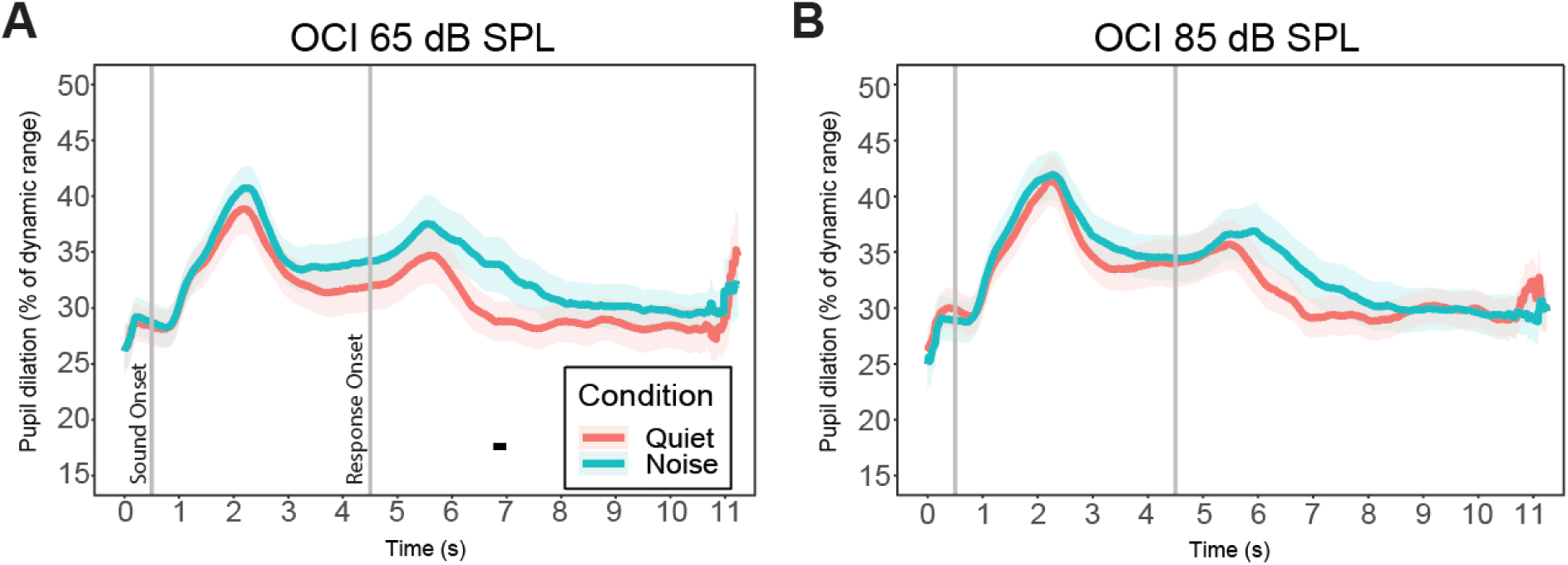
Pupillometry data for older CI users. A) Mean pupil dilation as a percentage of dynamic range across the timescale of each trial of experimental task performed in quiet (orange) and in the presence of noise (cyan) at 65 dB SPL for OCI listeners. B) Same as A but at 85 dB SPL. Shaded error bars indicate ± 1 standard error. Black bars indicate significantly different pupil size between quiet and noise conditions at the p < 0.05 level across the timescale shown.

### Listening effort is predicted by sound level, but effort does not predict behavioral performance

As observed from our pupillometry data, listening effort does in fact increase when stimuli are presented in noise at 85 dB SPL. To confirm whether listening effort is directly related to sound level, we ran a linear mixed effects model with our factors to predict pupil size. The final optimal model found a significant main effect of Level (β = 0.051, SE = 0.0155, p = 0.001) predicting the effort expended measured by the difference between pupil size in the noise and the quiet condition. Finally, we wanted to ask whether effort can predict the behavioral performance. However, the model did not show a main effect of Effort or any interactions with Effort, but the optimal model did demonstrate a significant main effect of Level (β = 0.0365, SE = 0.0055, p < 0.001) and an interaction of Group x Level (β = 0.0296, SE = 0.0055, p < 0.001), corresponding to our initial behavioral findings.

## Discussion Summary of results

The purpose of the study was to investigate the phenomenon of rollover as it relates to speech in noise performance among different age groups and hearing profiles. Here we found that behavioral performance and listening effort were modulated by sound level and the effect of rollover was mainly present when background noise was present. We compared this across subject groups and rollover was exacerbated by age and hearing loss in the acoustic listeners. CI users demonstrated no rollover effect in their behavioral performance or listening effort and this will be further discussed below.

### The effect of spectral and temporal deficits on rollover

One of our initial hypotheses was that rollover was correlated with a temporal processing deficit. Thus, we tested temporal contrast and non-temporal contrast word pairs. The temporal contrast word pairs directly tested this hypothesis while the non-temporal contrasts included phoneme differences along several dimensions including spectral aspects. Our results revealed a significant main effect of “Temporality” where there were differences in the behavioral performance functions between temporal and non-temporal word pair performances (Fig. 2). Previous studies revealed that high-pass filtering speech resulted in worse perception at high sound intensities than low-pass filtering speech (Molis and Summers, 2003), suggesting that high-frequency speech information may be more susceptible to rollover, leading to spectral hypotheses of rollover. These results correspond with our findings that spectral aspects of speech also do contribute to rollover to an extent. Future investigations are needed to understand the nuances of specific frequency-related changes across different phonemes.

Furthermore, when natural speech components, such as Consonant-Vowel (CV) combinations, are presented in noise, it was shown that rollover is affected by both spectral and temporal aspects. This includes features such as duration and nasality of a particular consonant, as well as place of articulation (Hornsby et al., 2005). These results correspond with our findings that these factors combined contribute to rollover as seen in both temporal and non-temporal word pair performance functions (Fig. 2). It is particularly interesting that the decreased spectral resolution from our CI users compounded with the effect of decreased temporal resolution of aging led to our CI group demonstrating relatively similar performance across levels in both temporal and non-temporal word pairs. It is known CI users make greater use of temporal cues because of the spectral degradation (Xu et al., 2005; Xu and Zheng, 2007), which may explain why performance was slightly better in the non-temporal word pairs than the temporal word pairs. Our result with CI user performance suggests two possibilities as follows. One possibility is that the spectrotemporal information lost from sounds passing through a CI also led to the loss of the spectrotemporal contributions to rollover. The information loss also led to decreased performance overall, especially in the noise conditions. The other possibility is that CI users may experience rollover differently than acoustic listeners, which may be affected by CI preprocessing strategies such as automatic gain control (AGC). The next steps would be to use vocoded speech in noise are needed to investigate in further detail how acoustic listeners and CI users experience rollover differently.

### Relating physiology to behavior across sound levels

One key finding of our study was that sound level is a main predictor of both behavior and physiology (pupillometry). Previous research confirms the level dependence of rollover, wherein NH acoustic listeners’ speech recognition performance decreased for levels above 75 dB SPL (Molis and Summers, 2003). Likewise, HI listeners have shown similar reductions in speech recognition for stimuli 87.5-100 dB SPL (Summers and Cord, 2007). However, these studies do not investigate across acoustic listening groups or CI users, and do not account for aging effects. The lack of research in this area may be attributed to the floor and ceiling effects of behavioral performance depending on the specific listening group that was tested. In addition, previous studies used speech-in-noise tests as a measure of listening effort in both in NH and HI listeners (Kramer et al., 1997; Zekveld et al., 2010, 2011; Koelewijn et al., 2012). Ohlenforst et al. (2017) demonstrated that HI listeners showed an increased pupil diameter to reach similar speech understanding performance compared to NH listeners. This supports our findings that HI listeners expend more effort with more difficult listening conditions, such as one with speech in noise at high sound levels in rollover. Our results showed that OHI listeners not only expended more effort than NH listeners, but also sustained their pupil response throughout the trial in both the listening and response phases when assessing the relative pupil dilation increase from quiet to noise conditions at 85 dB SPL (Fig. 5B). There are however several issues which arise, such as audibility at low presentation levels (35 dB SPL) in our study where there was no pupil size differences between the quiet and noise conditions in OHI listeners. This was similarly true for OCI users where there were no differences even across all sound levels presented (Fig. 6 and Supplementary Fig. 1D). These results may indicate factors which contribute to listening effort in general, since the difficulty of the task may influence whether the participant expends effort between the two SNR conditions (Pichora-Fuller et al., 2016). However, the control of these extenuating factors is often difficult and future investigations should aim to specifically account for these additional factors.

The novel nature of this study was that there has not been a previous investigation which attempts to relate the physiological effects of listening effort to the behavioral performance across levels, as most listening effort studies have focused on listening to speech in noise at a particular “comfortable” level, which cannot reveal the rollover effects seen here. Finally, while our results revealed that effort was a not a significant predictor of behavioral performance, it is likely that the main effect of level found in both indicates that the two measures co-vary and that other factors beyond listening effort contribute to the rollover effect found in the behavioral performance. It is also likely that listening effort has a relatively small range of changes compared to the group level effects observed in behavioral performance. Future studies should focus on elucidating additional contributions to rollover and how this differs across listening groups.

### CI users and level effects on speech-in-noise understanding

It is well known that there is great variability among CI users for speech recognition in noise tasks (Holden et al., 2013; Goehring et al., 2021). The variability among this subject group may be attributed to a number of factors including the varied programming and audibility features, duration of deafness, and age of implantation for both devices. Current research reinforces that early access to auditory stimulation is associated with better speech perception (Zaltz et al., 2020). Other factors associated with device programming and features can vary according to the manufacturer and the desired stimulation rate associated with the individual’s hearing profile. Specifically, each manufacturer utilizes its own pre-processing strategies which includes features such as compression of high intensity sounds or noise reduction relevant to our study here. Of note, Cochlear applies AGC and automatic sensitivity control (ASC) features which are designed to activate in the presence of impact sounds and dampen the sound’s intensity. The stimuli used in this study were two words presented in rapid succession and the background noise in the noise conditions was 6-talker babble noise, which have different spectrotemporal properties than an impact sound. Therefore, we believe the features of AGC/ASC did not necessarily interfere with the correct presentation level of our stimuli. The behavioral performance across levels in CI participants did change with sound level, and it was only the pupillometry data which showed similarities between the quiet and noise conditions. This could indicate that listening effort plays a different role in speech processing in CI users and perhaps rollover is experienced differently from acoustic listeners, if any at all. Given the variability of both the participants and CI programming, further investigation is needed into how CI users process speech at higher sound levels, as well as at various SNRs.

### Implications for other sensory systems

While our investigation here demonstrated rollover as an example of how high input levels can be detrimental to perception in the auditory system, this is similarly true in other sensory systems such as the visual system. Even in healthy aging, there is a decline in visual acuity in their ability to detect high contrast small targets (Madden and Greene, 1987; Skeel et al., 2003) as well as visual contrast sensitivity where there is a steep decline for high spatial frequencies and that extends to all frequencies after about 60 years old (Cabeza et al., 2005). It has also been consistently shown that these changes are due to higher internal equivalent noises or lower calculation efficiencies in older adults. Using flicker adaptation, an exposure to high temporal contrast, recent studies have shown reduced contrast sensitivity with aging in the magnocellular pathway but not in the parvocellular pathway (Zhuang et al., 2015). The magnocellular pathway is known to have high sensitivity to very low contrast, and the neural response rate increases quickly with increasing contrast, but quickly saturates at relatively low contrast. This oversaturation may have an equivalence in the auditory system as the effects of aging lead to denervation of a subset of low-spontaneous rate auditory nerve fibers responsible for processing suprathreshold stimuli (McClaskey et al., 2022). It is possible that sensory systems share mechanisms in which presentation of suprathreshold stimuli or oversaturation of neural activity is key to understanding why they are detrimental to perception. Future investigations should therefore take advantage of the auditory system and focus on the neural basis of rollover. With this particular focus, one can determine the locus of where the breakdown in neural coding begins and how this is affected by aging and hearing loss. Finally, understanding these neural mechanisms would inform us on aging and multisensory integration as well as how to better develop treatments for specific visual and hearing disorders.

## Conclusions

The present study demonstrated that rollover is not limited to retrocochlear pathologies and is a general listening phenomenon which affects acoustic listeners to different extents due to age and hearing loss. Additionally, listening effort plays a role, at least in part, in rollover due to changes observed at increasingly high sound levels. Clinically, the results of this study have implications on the diagnosis and treatment of age-related hearing loss. With devices such as hearing aids, they need to be carefully considered when programming amplification levels, as rollover can be detrimental to speech understanding at high sound intensities. Finally, with devices which treat severe hearing loss such as the CI, the evidence of rollover or lack thereof in CI demonstrated in this study has additional implications for aided assessments of functional listening and presentation level selection.

## Acknowledgements

We would like to thank Milena Costantino and Rebecca Farrar for their assistance with data collection. Research reported in this publication was supported by the National Institute on Deafness and Other Communication Disorders of the National Institutes of Health under Award Number R01DC020316 (MJG). The content is solely the responsibility of the authors and does not necessarily represent the official views of the National Institutes of Health.

---

**Supplementary Figure 1.**
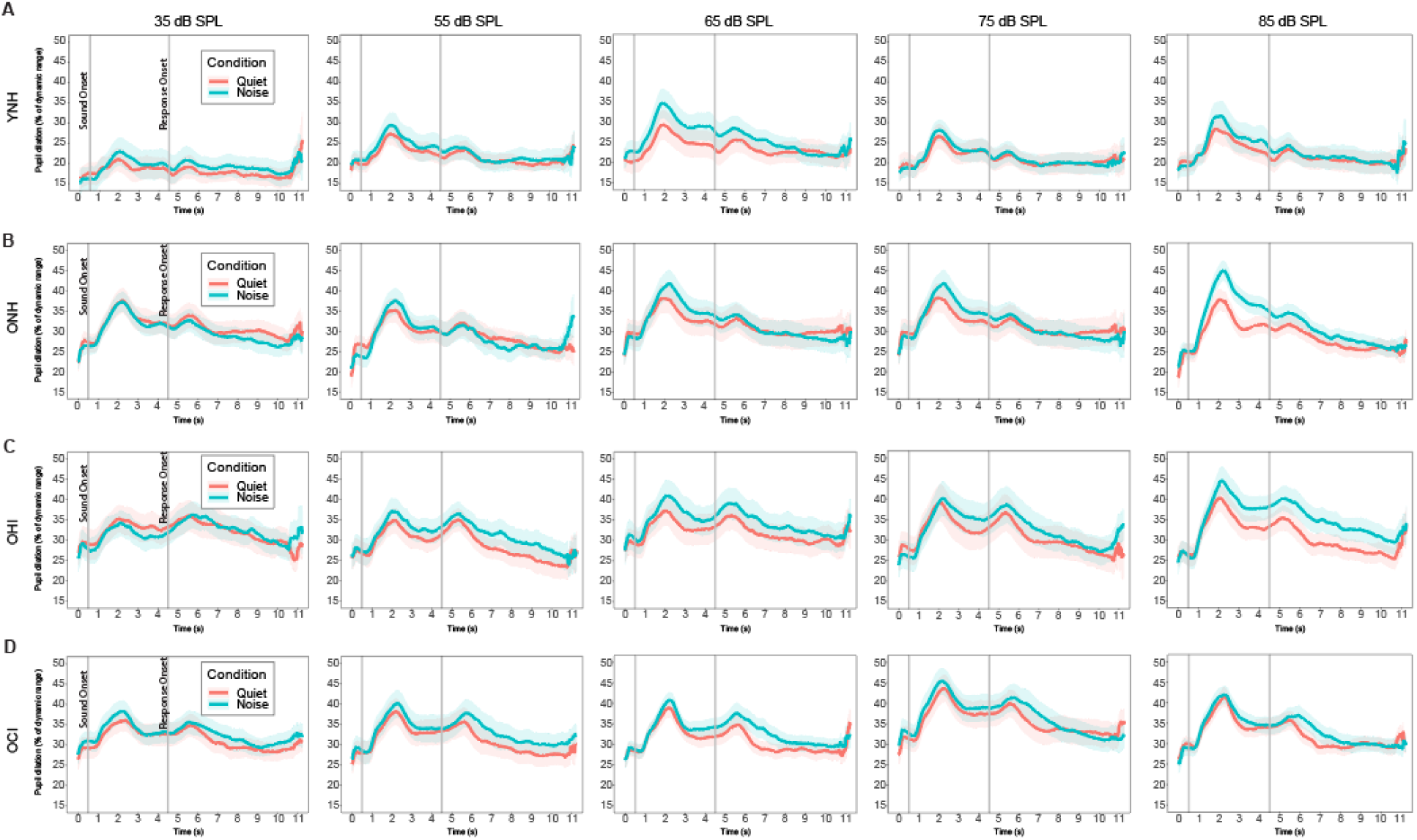
Pupillometry data for all groups. A) Mean pupil dilation as a percentage of dynamic range across the timescale of each trial of experimental task performed in quiet (orange) and in the presence of noise (cyan) at 35-85 dB SPL for YNH listeners. B, C, and D) Same as A but for ONH, OHI, and OCI groups respectively. Shaded error bars indicate ± 1 standard error.

**Supplementary Table 1.** Group comparisons of behavioral performance for each sound level in quiet condition. A) Independent samples t-test comparison results with Bonferroni adjusted p-values for behavioral performance scores in quiet condition at 35 dB SPL. B, C, D, and E) Same as A but for 55, 65, 75, and 85 dB SPL respectively. Significant p-values are bolded in each sub-table.

| **35 dB** | YNH | ONH | OHI | OCI |
| --- | --- | --- | --- | --- |
| YNH |  | 1.00 | **< 0.001** | **< 0.001** |
| ONH |  |  | **< 0.001** | **< 0.001** |
| OHI |  |  |  | 0.514 |
| OCI |  |  |  |  |

**B)**
| **55 dB** | YNH | ONH | OHI | OCI |
| --- | --- | --- | --- | --- |
| YNH |  | p = 0.47 | p = 1.00 | p = 0.866 |
| ONH |  |  | **p = 0.020** | **p = 0.003** |
| OHI |  |  |  | p = 1.00 |
| OCI |  |  |  |  |

**C)**
| **65 dB** | YNH | ONH | OHI | OCI |
| --- | --- | --- | --- | --- |
| YNH |  | p = 1.00 | p = 1.00 | p = 1.00 |
| ONH |  |  | p = 0.092 | p = 0.095 |
| OHI |  |  |  | p = 1.00 |
| OCI |  |  |  |  |

**D)**
| **75 dB** | YNH | ONH | OHI | OCI |
| --- | --- | --- | --- | --- |
| YNH |  | p = 1.00 | p = 1.00 | p = 0.307 |
| ONH |  |  | p = 1.00 | **p = 0.004** |
| OHI |  |  |  | p = 0.05 |
| OCI |  |  |  |  |

**Supplementary Table 1. Group comparisons of behavioral performance for each sound level in quiet condition.**
| **85 dB** | YNH | ONH | OHI | OCI |
| --- | --- | --- | --- | --- |
| YNH |  | p = 1.00 | p = 1.00 | p = 1.00 |
| ONH |  |  | p = 0.916 | p = 0.086 |
| OHI |  |  |  | p = 1.00 |
| OCI |  |  |  |  |

**Supplementary Table 2.**
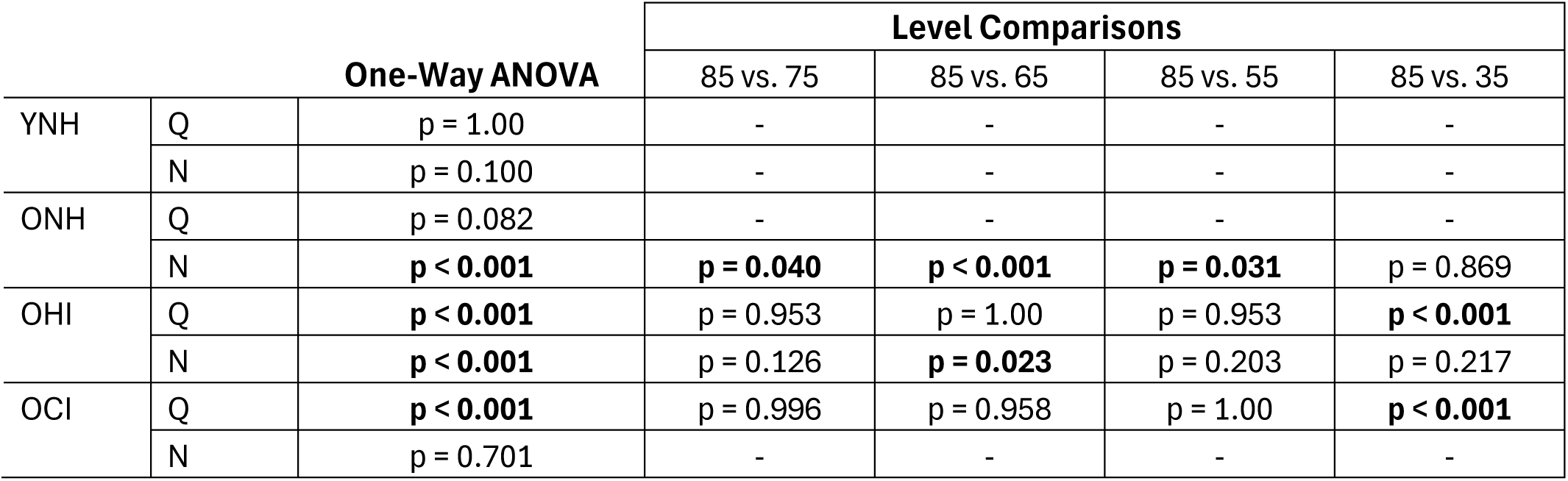
Individual level comparisons of behavioral performance against the highest sound intensity (85 dB SPL) for each group in quiet and noise conditions. One-way ANOVAs were performed in each group to determine if there were significant differences in behavioral performance across levels. If there were significant differences, a post-hoc multiple comparisons test with Bonferroni corrections was performed for 85 dB SPL against the other levels to determine if rollover was present. Note that the significant differences found between 85 dB vs. 35 dB SPL in quiet condition were due to lower performance at 35 dB SPL rather than rollover. Significant p-values are bolded in the table.

## Notes

### Competing Interest Statement

The authors have declared no competing interest.

